# Acute exposure to nicotine vapor causes short-term increases in impulsive choice in rats

**DOI:** 10.1101/842005

**Authors:** R.J. Flores, F.Z. Alshbool, P. Giner, L.E. O’Dell, I.A. Mendez

## Abstract

**Background:** Previous studies have shown that exposure to nicotine smoke increases impulsivity. Surprisingly, research investigating the effects of electronic cigarette nicotine vapor exposure on impulsivity has not been conducted. Therefore, the present study examined the effects of nicotine vapor exposure on impulsive choice.

**Methods:** Twenty-four adult male rats were trained in the delay discounting task to choose between small immediate food rewards or larger food rewards with delayed deliveries. After 24 days of training in the delay discounting task, rats were passively exposed to vapor containing either 0, 12, or 24 mg/mL of nicotine for 10 days. To monitor exposure to nicotine, serum cotinine levels were assessed on exposure days 1, 5, and 10 using enzyme-linked immunosorbent assay (ELISA). Following vapor exposure, rats were retrained in the delay discounting task until stable performance was achieved, and the effects of nicotine vapor exposure on choice preference were assessed.

**Results:** Rats that were exposed to 12 and 24 mg/mL nicotine vapor displayed higher serum cotinine levels, relative to those exposed to 0 mg/mL nicotine vapor. There were no differences in impulsive choice between any of the vapor groups when tested 15-21 days after exposure. However, increases in impulsive choice were observed when testing immediately following exposure to 24 mg/mL nicotine vapor, relative to immediately following exposure to 0 mg/mL nicotine vapor.

**Conclusions:** Findings suggest that while exposure to nicotine vapor may not cause long-term changes in decision making, it can cause short-term increases in impulsive choice, an effect that can have negative social and health consequences.

## 1. Introduction

The effects of smoking on the brain and behavior have been well documented. Both clinical and preclinical research has shown that repeated exposure to nicotine causes aberrant neurological and behavioral processes (Ascioglu, Dolu, Golgeli, Suer, & Ozesmi, 2004; Dani & Harris, 2005; Mansvleder & McGehee, 2002; Yuan, Cross, Loughlin, & Leslie, 2015). Research investigating the effects of nicotine vapor inhalation from electronic cigarettes (e-cigarettes) is beginning to demonstrate that consumption of nicotine using this novel nicotine delivery system can have detrimental effects on human health that are both similar and new when compared to traditional smoking (Callahan-Lyon, 2014; Chun, Moazed, Calfee, Matthay, & Gotts, 2017). However, pre-clinical studies linking nicotine vapor consumption from e-cigarettes to changes in the brain and behavior are limited (Javadi-Paydar, Kerr, Harvey, Cole, & Taffe, 2019; Smith et al., 2015). This is concerning, as e-cigarette vaping in adolescents has now surpassed conventional cigarette smoking, with use increasing from 1.5% in 2011 to 20.8% in 2018 (Gentzke et al., 2019). The limited findings that are available thus far suggest that while nicotine vapor may contain fewer toxic additives than nicotine smoke, higher administration rates and chemical constituents associated with e-cigarettes can enhance nicotine’s addiction-related behavioral and biological effects (Gilpin et al., 2014; Hall et al., 2014). Additionally, the high concentrations of nicotine found in e-cigarette solutions, along with increased frequency of use, may severely impair respiratory function and increase the likelihood of future conventional cigarette use (Chun et al., 2017; Lozano et al., 2017).

One effect of traditional cigarette use that has been well investigated is the relationship between smoking and impulsivity (Bickel, Odum, & Madden, 1999; Chivers, Hand, Priest, & Higgins, 2016; Mitchell, 2004). Many preclinical studies have investigated the effects of nicotine injections on impulsive choice in rodents; however, this research has produced mixed findings (Anderson & Diller, 2010; Dallery & Locey, 2005; I A Mendez, Gilbert, Bizon, & Setlow, 2012). Alternatively, no studies examining the effects of nicotine vapor exposure from e-cigarettes on impulsive choice have been conducted.

There is a general lack of research on the effects of nicotine vapor on the brain and behavior and this may be due, in part, to the limited number of validated nicotine vapor delivery systems designed for animal research. The present study used a relatively novel nicotine vapor delivery system (Qasim et al., 2018) to investigate the effects of nicotine vapor exposure on impulsive choice, in rats transitioning from adolescence to adulthood.

## 2. Methods

### Subjects

Male Sprague-Dawley rats (N=24) were obtained at 3 weeks of age from an out-bred stock of animals (Envigo, Inc., Indianapolis, IN). The rats were paired-housed in a humidity- and temperature-controlled (22°C) vivarium on a reverse 12-hour light/dark cycle (lights off at 6:00 am and on at 6:00 pm) with ad libitum access to food and water, except when noted below. All animal procedures were performed in accordance with the University of Texas at El Paso animal care committee’s regulations.

### Experimental design

Prior to beginning the experiment, the rats were handled in the vivarium for 2 consecutive days. The experiment schedule (Table 1) began with shaping procedures for the delay-discounting task (as a measure of impulsive choice), which included magazine training, lever press training for a grain food pellet (left and right lever, counter balanced), and nose-poke training, in that order, until a criteria 30 responses in 1 hr was met at each stage. Rats were then trained daily in the delay discounting task for 24 non-treatment days. To investigate the effects of nicotine vapor exposure during young adulthood, separate groups of 10 week old rats were exposed to vaporized nicotine liquid (0, 12, or 24 mg/mL nicotine) for 10 days, with 2 days off between exposure days 5 and 6 (Sengupta, 2013). To verify the delivery of nicotine in rats, blood collection procedures were conducted immediately following vapor exposure on days 1, 5, and 10, and blood serum levels of its metabolite cotinine were assessed. Following nicotine vapor exposure procedures, the rats resumed non-treatment training in the delay discounting task until criterion for stable responding was achieved across 6 consecutive days, which occurred on non-treatment delay discounting days 35-40 (DD 35-40; see Simon, Mendez, & Setlow, 2007 for more information). To examine the effects of acute nicotine vapor exposure on impulsive-choice, all 24 animals were pooled and tested in the delay-discounting task immediately following 0 mg/mL nicotine vapor exposure on treatment delay discounting test day 1 (TD1) and 24 mg/mL nicotine vapor exposure on treatment delay discounting day 2 (TD2). To assess potential lingering effects of 24 mg/mL nicotine vapor exposure on impulsive choice, one final non-treatment delay discounting day (DD 41) was conducted 24 hours after TD2.

**Table 1.** Experiment Schedule for Training, Testing, and Vapor Exposure

| Group | Training<br>Phase I<br>DD 1-24 | 10 Days<br>Vapor<br>Exposure | Training<br>Phase II<br>DD 25-34 | Stability/Long-Term Test<br>DD 35-40 | Short-Term<br>Test<br>TD 1 | Short-Term<br>Test<br>TD 2 | 24hrs post<br>Exposure<br>DD 41 |
| --- | --- | --- | --- | --- | --- | --- | --- |
| 1 | No<br>Exposure | 0 mg/ml<br>Nicotine | No<br>Exposure | No<br>Exposure | 0 mg/ml<br>Nicotine | 24 mg/ml<br>Nicotine | No<br>Exposure |
| 2 | No<br>Exposure | 12 mg/ml<br>Nicotine | No<br>Exposure | No<br>Exposure | 0 mg/ml<br>Nicotine | 24 mg/ml<br>Nicotine | No<br>Exposure |
| 3 | No<br>Exposure | 24 mg/ml<br>Nicotine | No<br>Exposure | No<br>Exposure | 0 mg/ml<br>Nicotine | 24 mg/ml<br>Nicotine | No<br>Exposure |

\*DD = Non-Treatment Delay Discounting Day, TD = Treatment Delay Discounting Day

### Nicotine Vapor Exposure

To model e-cigarette nicotine vapor exposure, rats received passive nicotine vapor exposure 5 days a week (Monday-Friday) using the Four Chamber Benchtop Passive E-Vape Inhalation System (La Jolla Alcohol Research Inc., La Jolla, CA; Qasim et al., 2018). This system consisted of four sealed chambers (interior dimension of 14.5” L x 10.5” W x 9.0” H), each with two valve ports. One valve port was connected to a small vacuum that controlled the airflow in the chamber at 0.6 L per minute. The vacuum outlet was connected to a Whatman HEPA filter (Millipore Sigma, Darmstadt, Germany) and onto a house exhaust that safely removed the nicotine from the chambers and outside the testing room. The other valve port was connected via PVC tubing to a modified 4.9 volt TFV4 minitank (Smok Inc, Shenzhen, China), where the nicotine was heated. The minitanks are also linked to a control box that allows for controlled heating of nicotine e-cigarette liquid (e-liquid).

The e-liquid nicotine concentrations and exposure protocols with rodents were selected to model daily human nicotine consumption rates and serum cotinine levels from e-cigarette use (Etter, 2016; Flouris et al., 2013; Qasim et al., 2018). At 10 weeks of age (the approximate period when rats are transitioning form adolescence to adulthood) rats were exposed to either 0, 12, or 24 mg/mL nicotine vapor for 10 days, with 2 days off between days 5 and 6 of vapor exposure. Two rats were exposed per chamber (cagemates) for a total of 8 rats per exposure session, with each nicotine vapor group exposed in separate sessions each day. Since all 4 chambers were connected to 1 minitank, all 8 rats in a given session had to be exposed to the same concentration of nicotine. To minimize the possibility of higher concentrations contaminating lower concentrations, 8 rats were exposed to the 0 mg/mL nicotine concentration in the first session, 8 rats were exposed to the 12 mg/mL nicotine concertation in the second session, and 8 rats were exposed to the 24 mg/mL concentration in the third session. In an attempt to avoid cross contamination, separate tubing and minitanks were used for each concentration of nicotine e-liquid and chambers were thoroughly cleaned after every exposure. Vapor Chef brand flavorless nicotine e-liquids (a common choice among teens) containing 0, 12, or 24 mg/mL nicotine in 50/50 vegetable glycerin/propylene glycol vehicle were used in this study. The rats were exposed to vapor in daily sessions consisting of 4 cycles, with 5-minute inter-cycle-intervals. For each cycle, the 4.9 volt minitanks heated the nicotine e-liquid to 400 degrees Fahrenheit for a 3 second puff duration. The resulting vapor puff was delivered to the rats every 2 minutes and this process continued until the rats received 10 vapor puff deliveries per cycle, for a total of 40 puffs per daily session.

### Assessment of cotinine levels

Immediately following vapor exposure procedures on exposure days 1, 5, and 10, the rats were briefly anesthetized using an isoflurane/oxygen mixture (1-3% isoflurane) and received a nick on the end of the tail with a sterile scalpel blade. Blood was collected from the tail in sterile 1 mL Eppendorf tubes and placed on ice. The blood was then centrifuged for 15 min at 5000 x g at 4°C. Serum was extracted and stored in aliquots at −80°C until analyzed via Enzyme-linked immunosorbent assay (ELISA) procedures for cotinine analysis according to manufacturer instructions (Cal Biotech Inc, El Cajon, CA). The standards ranged from 0 to 100 ng/mL of cotinine and were included in every assay. Samples were diluted 1:4 in 1X PBS with 1% – 2% BSA (Fraction V grade) based on a sample dilution pilot study revealing that a 4x dilution permitted the detection of cotinine within the standard curve range. Samples were loaded onto a 96 well-plate and read at 450 nm wavelength using a Spectra Maxplus spectrophotometer (Molecular Devices Inc, Sunnyvale, CA). Obtained serum cotinine values were then multiplied by 4 to correct for our 1:4 dilution and determine the final serum cotinine values.

### Delay Discounting Task

The present study utilized 8 standard operant chambers and procedures previously described in our work (I.A. Mendez et al., 2010; Simon et al., 2007). Operant chambers (interior dimension of 12.0” L x 9.5” W x 11.5” H) were placed inside a sound-attenuating PVC shell (interior dimensions: 14.0” L x 22.0” W x 15.0” H; MED associates, St. Albans, VT) and had a 1.12 W house light that was illuminated throughout training and testing. Operant sessions were conducted using 2 retractable levers that extend 2.5 cm into the chamber. A pellet dispenser located outside of the operant chamber, but inside the chamber shell, was used to deliver 45-mg food pellets into a metal trough located within the chamber.

Five days prior to beginning the delay discounting procedures, rats were food restricted to 90% of their free-feeding bodyweight. Feeding was only allowed for 2-hours per day, after behavioral testing was completed for the day. The task began with rats receiving two 45 min sessions of magazine training consisting of 16 to 22 deliveries of a 45 mg food pellet with an average delivery interval of 2 minutes. This was followed by 4 consecutive 1 hr sessions of lever press training where rats were presented with a single lever (2 days with left lever and 2 days with right lever) and allowed to press for a food pellet under a fixed ratio 1 schedule of reinforcement. Upon the completion of lever press training (30 presses within the 1 hr sessions), the rats were trained to nose poke into an illuminated food trough, after which the 1.12 W trough light was turned off, and a single lever (left or right) was presented. When the rats pressed on the presented lever, two food pellets were delivered and the lever was retracted. All left and right lever press assignments and extensions were randomly assigned and counterbalanced during these sessions.

The delay discounting task was conducted for 41 non-treatment days and 2 treatment days (acute nicotine vapor treatment days), which occurred between non-treatment days 40 and 41. Each discounting session consisted of 5 blocks, each with 4 forced choice trials (no lever option) followed by 6 free choice trials (two lever options). Individual trials began with the presentation of a light cue (10s food cup-light), which was extinguished when the rat entered the food trough, initiating lever insertion. Pressing one of the two levers (left or right of the food cup, randomly assigned and counterbalanced) resulted in the immediate delivery of a small reward (one grain food pellet), whereas pressing the other lever resulted in the delayed delivery of a large reward (4 food pellets) with a determined delay for delivery. The delay to the large reward delivery decreased across session blocks (0, 4, 8, 16, 32 seconds). Once either lever was pressed, both levers were retracted until the next trial. Performance in this task was assessed once stable performance was achieved in the delay discounting task (beginning on non-drug training day 35). Stable performance was defined as a significant effect of delay block with no significant effect of training day, across 6 consecutive training days (see (Simon et al., 2007) for more information), which occurred on training days 35 through 40.

### Statistical analysis

To examine short-and long-term effects of nicotine exposure on delay discounting, choice preference in delayed reward was compared using a mixed two-way ANOVA with group (0, 12, or 24 mg/mL nicotine) as a between subjects factor and delay block or training day as within subject factors. Independent samples t-test and Tukeys HSD were used for post-hoc analyses, while partial eta squared and Cohen’s d were used to determine effect sizes.

## 3. Results

### Training in the Delay Discounting Task

Rats were trained in the delay discounting task for 24 days before beginning young adulthood nicotine vapor exposure. Analysis of nose-poke and lever press omissions using One way ANOVA showed that there were no significant differences between the nicotine vapor exposure groups on the 5 non-treatment delay discounting days preceding vapor exposure, all training days after vapor exposure, or acute vapor exposure test days (*F_s_* < 2.39, *p_s_* > 0.12). A mixed model ANOVA revealed no significant main effects of, or interactions with, choice preference across the 5 non-treatment delay discounting days prior to nicotine vapor exposure (training days 20-24; *F_s_* <1.39, *p_s_>0.08).* Following vapor exposure, rats continued to train in the delay discounting task for an additional 10 days before stability in task performance was achieved across all rats. No significant main effects or interactions with nicotine treatment group were observed during these first 10 post-treatment delay discounting days (non-treatment delay discounting days 25-34, *F_s_*<1.12, *p_s_*>0.24), suggesting that nicotine vapor exposure does not affect learning in the delay discounting task. Criteria for stable responding in the delay discounting task was not met until non-treatment delay discounting days 35-40 (Figure 1), during which a significant main effect of block was observed (*F*_(4,80)_=81.91, *p*<0.001) without a significant main effect of day (*F*_(5,100)_=0.79, *p*=0.56).

**Figure 1.**
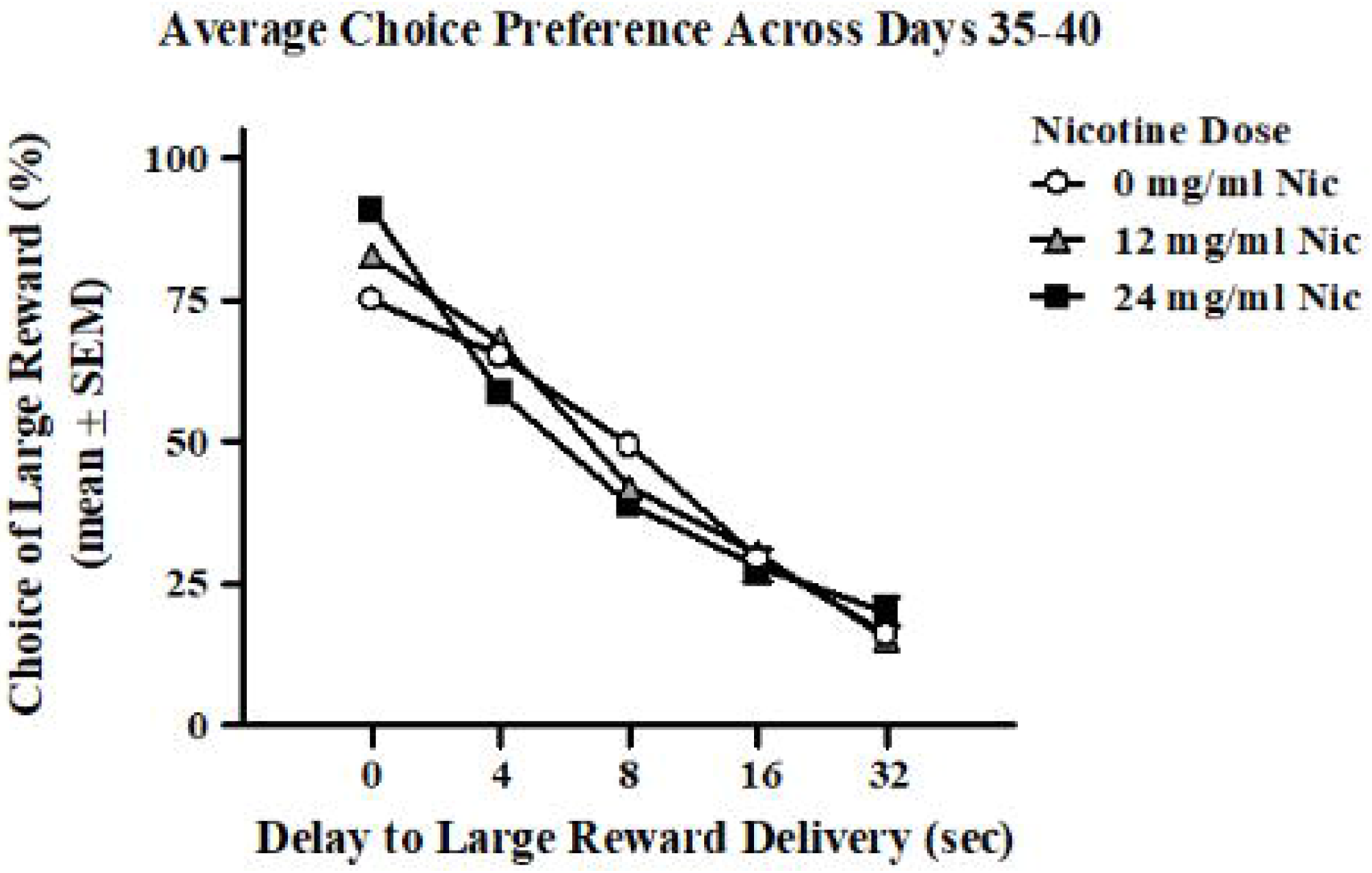
Long-term effects of nicotine vapor exposure on average choice preference across 6 days of stable responding. Rats showed no significant long-term effects on impulsive choice averaged across 6 days of stable responding (Non-Treatment Delay Discounting Days 35-40) when tested 15 days following exposure to 10 days of 0 (0 mg/mL nic, white circles), 12 (12 mg/mL nic, grey triangles), or 24 (24 mg/mL nic, black squares) mg/mL of nicotine vapor.

### Serum Cotinine

To confirm systemic nicotine delivery during the passive vapor exposure, blood cotinine levels were collected approximately 20 minutes after nicotine vapor exposure, on vapor exposure days 1, 5, and 10 (Figure 2). A mixed model ANOVA with nicotine vapor group as a between subject factor and blood collection day as a within subject factor revealed a significant main effect of nicotine vapor group (*F*_(2,21)_=81.10, *p*<0.001, *η_p_*^2^ =0.89) and blood collection day (*F*_(2,42)_=21.35, *p*<0.001, *η_p_*^2^ =0.51), as well as a significant interaction of nicotine vapor group and day (*F*_(4,42)_=6.51, *p*<0.01, *η_p_*^2^=0.38). One-way ANOVA revealed significant main effects of nicotine vapor exposure on days 1 (*F*_(2)_=17.31, *p*<0.001, *η_p_*^2^=0.62), 5 (*F*_(2)_=34.27, *p*<0.001, *η_p_*^2^ =0.77), and 10 (*F*_(2)_=57.56, *p*<0.001, *η_p_*^2^=0.85). Tukey HSD post-hoc analysis comparing cotinine levels between individual treatment groups at each cotinine analysis day reveal a significant difference between the 0 mg/mL nicotine group and both the 12 and 24 mg/mL nicotine groups, at all 3 time points (*p_s_*<0.01, Cohen’s d>2.83). No significant difference between the 12 mg/mL and 24 mg/mL nicotine treatment groups were observed during any of the 3 cotinine analysis time points (*p_s_*>0.13). Together, these findings suggest that passive exposure to 12 and 24 mg/mL nicotine vapor similarly increases nicotine delivery, relative to passive exposure to 0 mg/mL nicotine vapor.

**Figure 2.**
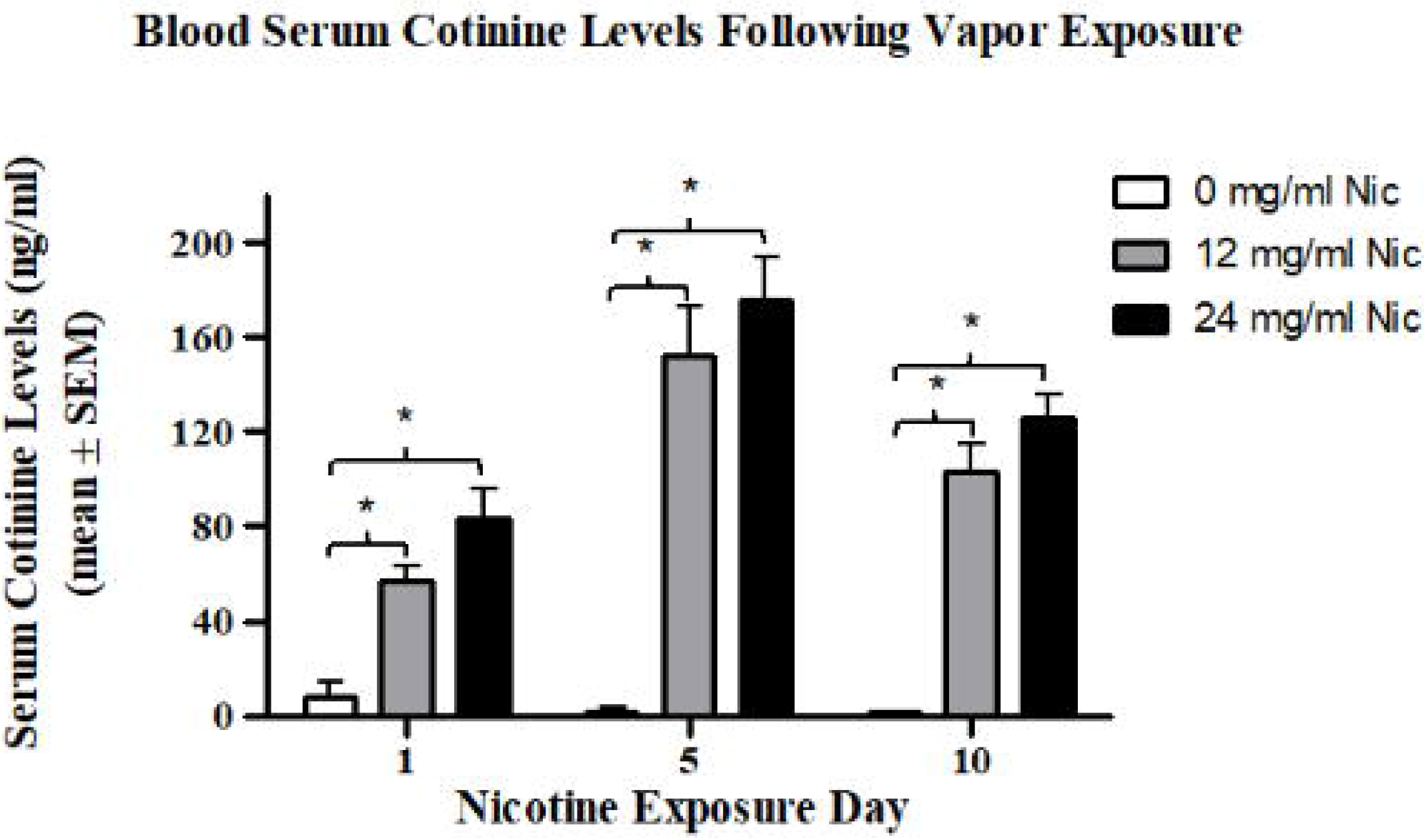
Blood serum cotinine levels of rats that were exposed to nicotine or vehicle vapor on exposure days 1, 5, and 10. The rats that were exposed to the 12 mg/mL (grey bars) and 24 mg/mL (black bars) nicotine vapor displayed an increase in serum cotinine levels when compared to the rats that were exposed to 0 mg/mL (white bars) nicotine vapor (ps<0.01). No significant differences were observed between the 12mg/mL and the 24 mg/mL nicotine treatment groups.). Data are presented as mean ± S.E.M. Asterisk (*) indicate a significant difference between the 0 mg/mL and 12 or 24 mg/mL nicotine vapor treatment groups. Critical p value is 0.05.

### Short- and Long-Term Effects of Nicotine Vapor Exposure on Impulsive Choice

To assess the long-term effects of 10 days of nicotine vapor exposure on choice preference in the delay discounting task, performance on the delay discounting task was compared between the 10-day nicotine vapor treatment groups on each of the 6 training days in which rats showed stable responding (DD 35-40), as well as the average performance across these 6 days (Figure 1). As described above in our statistical analysis of stability in the delay discounting task, a significant main effect of block was observed across non-treatment days 35-40 (*F*_(4,80)_=81.91, *p*<0.001). Repeated measures ANOVA were ran on each of these stability days and revealed a marginally significant interaction between 10-day nicotine vapor exposure day and discounting block on training day 37 (*F*_(8,84)_=2.02, *p*=0.05, *η_p_*^2^=0.16). No other significant interactions or main effects of nicotine vapor exposure were observed on any of the 6 individual non-treatment delay discounting days or the average of these 6 days (*F_s_*<1.58, *p_s_*>0.14). Post-hoc analysis were used to investigate the interaction observed on training day 37 and no significant differences between nicotine vapor groups were observed on any of the discounting blocks (*p_s_*>0.37). Together, these findings suggest that 10 days of 12 or 24 mg/mL nicotine vapor exposure does not have significant long-term term effects on impulsive choice.

Following analysis of the long-term effects of nicotine vapor exposure across 6 days of stable responding, the short-term effects of passive nicotine vapor exposure were assessed in each 10-day treatment group immediately following exposure to 0 mg/mL (TD1) and 24 mg/mL (TD2) nicotine vapor. No significant differences were observed in any 10-day nicotine vapor treatment group when using repeated measures ANOVA to compare choice preference on non-treatment day 40 to choice preference immediately following exposure to 0 mg/mL nicotine (*F_s_*<0.38, *p_s_*>0.56). Repeated measures ANOVA comparing choice preference immediately following 0 mg/mL nicotine vapor exposure to choice preference immediately following 24 mg/mL nicotine vapor exposure revealed a significant main effect of treatment day (*F*_(1,7)_=6.57, *p*<0.05, *η_p_*^2^=0.48) and block (*F*_(4,28)_=22.63, *p*=0.001, *η*_p_^2^=0.76) in the group that previously received 0 mg/mL nicotine vapor for 10 days. No interaction of day and block (*F*_(4,28)_=0.31, *p*=0.87) was observed in this group. This finding suggests that acute nicotine exposure during young adulthood can cause immediate increases impulsive choice (Figure 3). Post-hoc analysis further revealed that immediate increases in impulsive choice in this group following exposure to the 24 mg/mL nicotine vapor were driven by significant increases in impulsive choice during block 4 (16 second delay) of the delay discounting task (*t*_(7)_=3.21, *p*<0.05, Cohen’s d=0.71).

**Figure 3.**
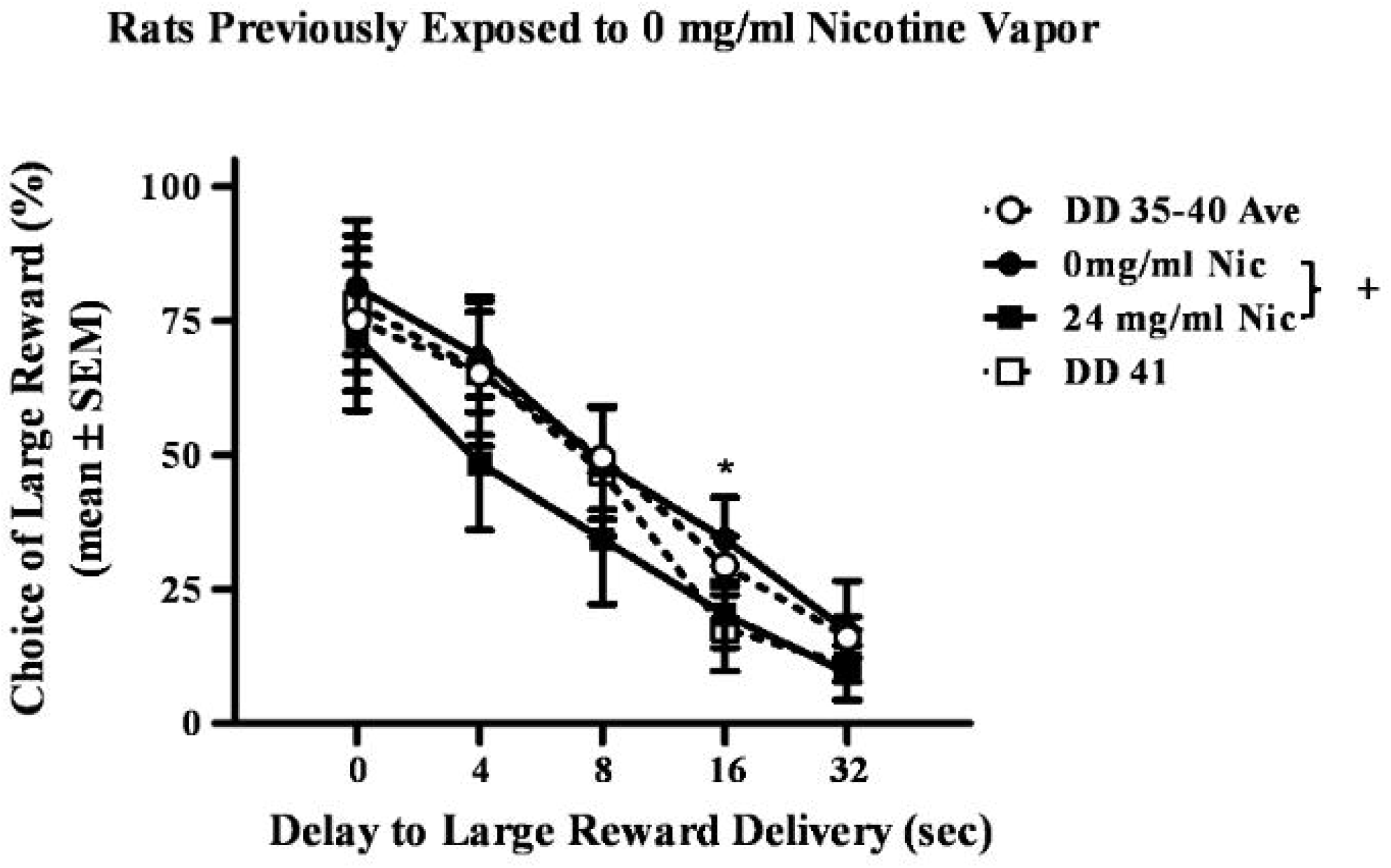
Short-term effects of nicotine vapor exposure on choice preference in rats previously treated with 10 days of 0 mg/mL nicotine. Rats previously exposed to 0 mg/mL nicotine vapor demonstrated increased impulsive choice (decreased choice of the large delayed reward). Data shown include choice preference averaged across days 35-40 (DD 35-40 Ave, white circle), immediately after exposure to 0 mg/mL nicotine vapor (0 mg/mL nic, black circle), immediately after exposure to 24 mg/mL nicotine vapor (24 mg/mL nic, black square), and 24 hours after exposure to 24 mg/mL nicotine vapor (DD 41, white square). Data are presented as mean ± S.E.M. Plus sign (+) indicates a significant effect of treatment day with asterisk (*) indicating a significant difference on block between the 0 mg/mL and 24 mg/mL nicotine vapor treatment days. Critical p value is 0.05.

The effects of acute nicotine vapor exposure were also assessed in rats with a history of exposure to 12 and 24 mg/mL nicotine vapor. As seen with the group that received 0 mg/mL nicotine vapor, a repeated measures ANOVA revealed a significant main effect of block (*F_s_*>32.79, *p*<0.001, *η_p_*^2^=0.82), but no significant interaction of acute treatment day and block (*F_s_*<1.24, *p_s_*>0.32), in both the 12 and 24 mg/mL nicotine groups. Interestingly, there was a significant main effect of acute 24 mg/mL nicotine vapor treatment day in the group with previous passive exposure to 10 days of 12 mg/mL nicotine vapor (*F*_(1,7)_=8.33, *p*<0.05, *η_p_*^2^ =0.54, Figure 4), but not in the group with previous exposure to 10 days of 24 mg/mL nicotine vapor (*F*_(1,7)_=0.32, *p*<0.59, Figure 5). Post-hoc analyses were conducted to characterize the main effect of treatment day observed in the 12 mg/mL nicotine group. Post-hocs comparing treatment day effects at each delay block, revealed that exposure to the 24 mg/mL nicotine concentration increased choice for the small immediate reward during block 4 (16 second delay), relative to choice preference during block 4 following 0 mg/mL nicotine vapor exposure (*t*_(7)_=3.33, *p*<0.05, Cohen’s d=1.13).

**Figure 4.**
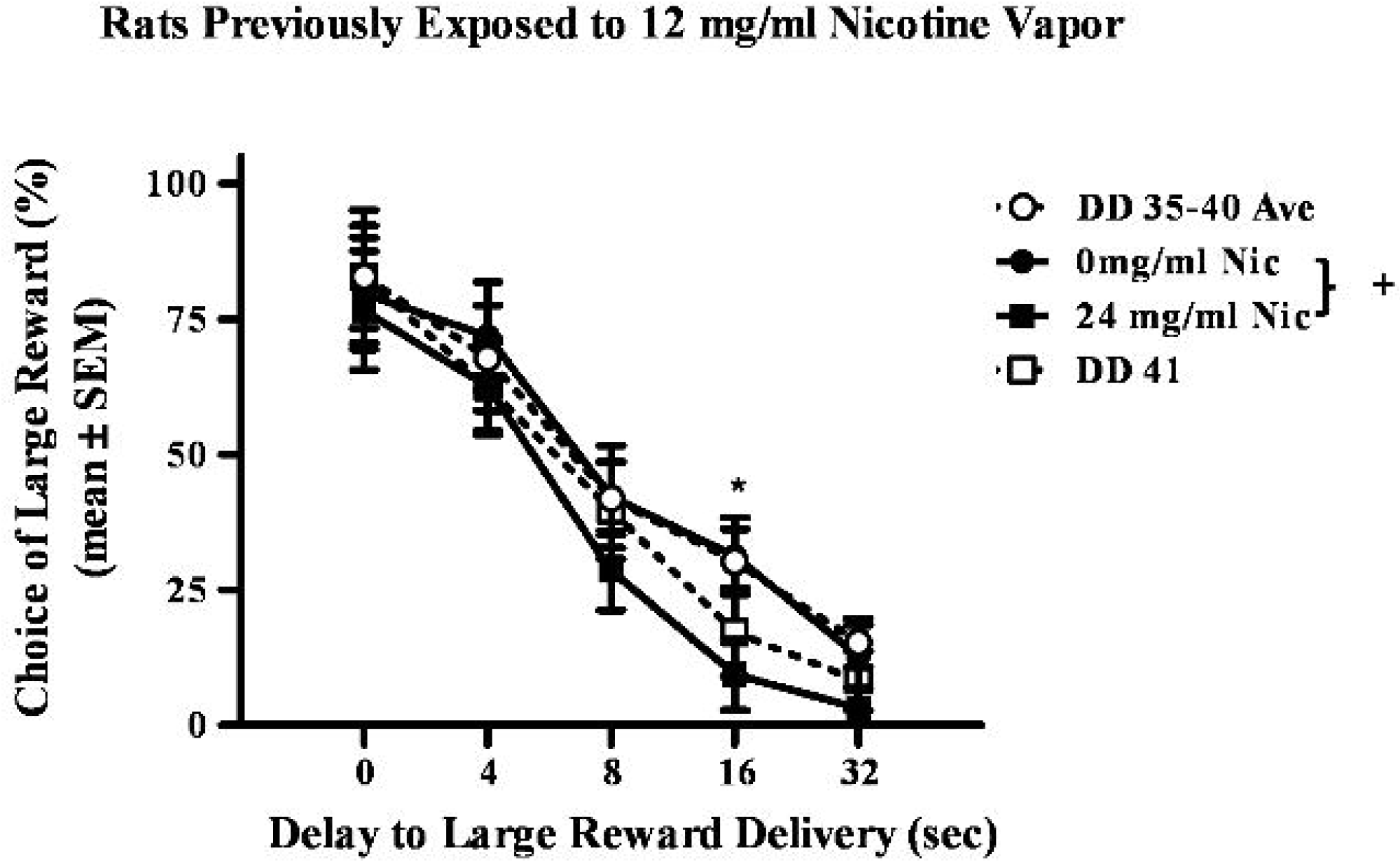
Short-term effects of nicotine vapor exposure on choice preference in rats previously treated with 10 days of 12 mg/mL nicotine. Rats previously exposed to 12 mg/mL nicotine vapor also demonstrated increased impulsive choice (decreased choice of the large delayed reward). Data shown include choice preference averaged across days 35-40 (DD 35-40 Ave, white circles), immediately after exposure to 0 mg/mL nicotine vapor (0 mg/mL nic, black circles), immediately after exposure to 24 mg/mL nicotine vapor (24 mg/mL nic, black squares), and 24 hours after exposure to 24 mg/mL nicotine vapor (DD 41, white circles). Data are presented as mean ± S.E.M. Plus sign (+) indicates a significant effect of treatment day with asterisk (*) indicating a significant difference on block between the 0 mg/mL and 24 mg/mL nicotine vapor treatment days. Critical p value is 0.05.

**Figure 5.**
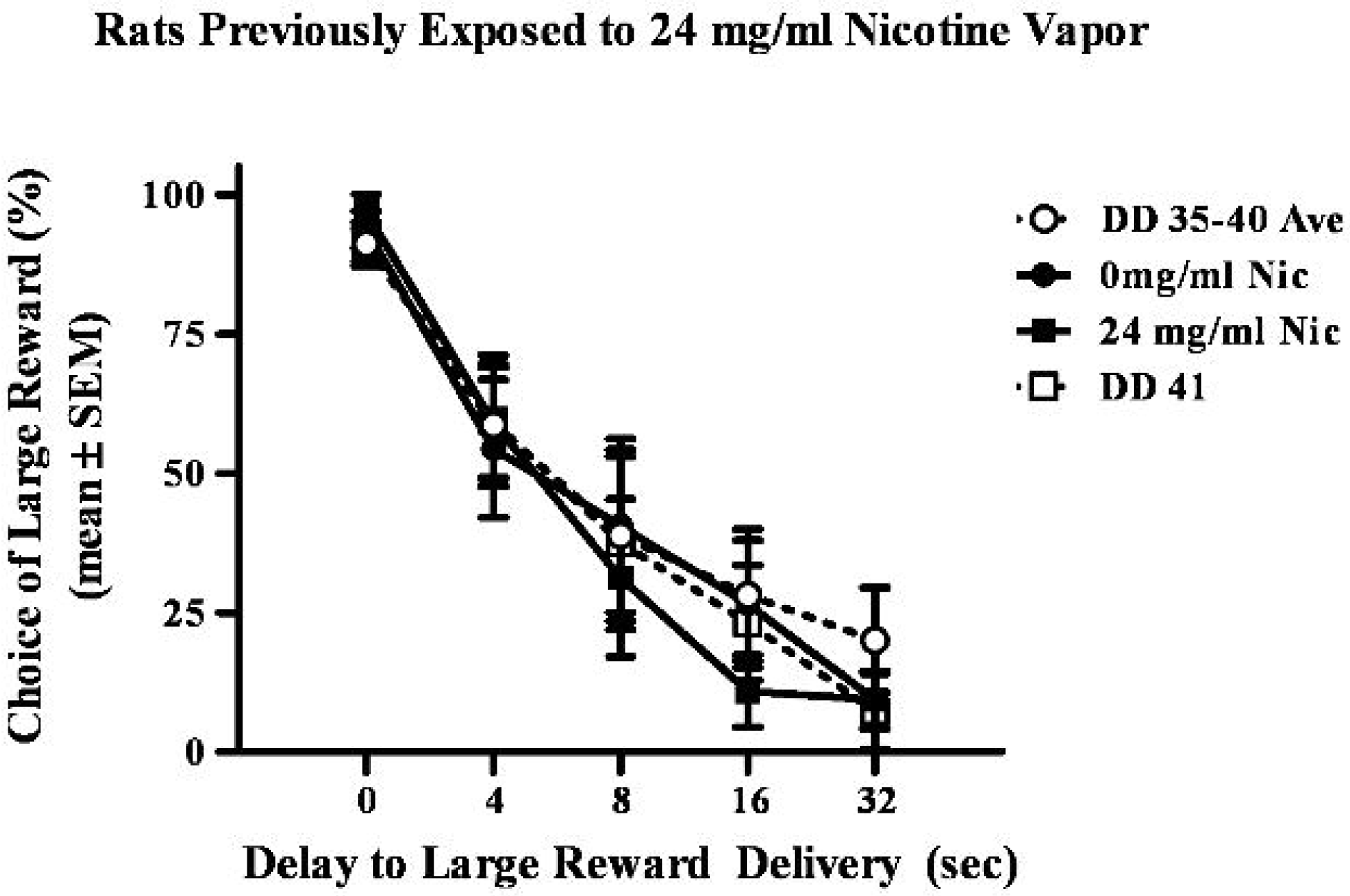
Short-term effects of nicotine vapor exposure on choice preference in rats previously treated with 10 days of 24 mg/mL nicotine. Rats previously exposed to 24 mg/mL nicotine vapor did not demonstrate increased impulsive choice (decreased choice of the large delayed reward). Data shown include choice preference averaged across days 35-40 (DD 35-40 Ave, white circles), immediately after exposure to 0 mg/mL nicotine vapor (0 mg/mL nic, black circles), immediately after exposure to 24 mg/mL nicotine vapor (24 mg/mL nic, black squares), and 24 hours after exposure to 24 mg/mL nicotine vapor (DD 41, white squares). Data are presented as mean ± S.E.M.

To assess the persistence of acute effects of nicotine vapor exposure on impulsive choice, a separate repeated measures ANOVA was ran comparing choice preference following acute exposure to 0 mg/mL nicotine vapor to choice preference 24 hours after acute exposure to 24 mg/mL nicotine vapor (DD 40). No significant effects of day were observed in any of the 10-day treatment groups when comparing choice preference 24 hours after exposure to 24 mg/mL to choice preference immediately following 0 mg/mL nicotine treatment (*F_s_*<2.42, *p_s_*>0.16). However, when comparing choice preference 24 hours after 24 mg/mL exposure to choice preference on training day 40, a notable near significant probability value (*F*_(1,7)_<4.84, *p*=0.06, *η_p_*^2^=0.41) was obtained in rats that previously received 10 days of 12 mg/mL nicotine vapor exposure. While findings were not significant, data suggests that acute vapor exposure to 24 mg/mL nicotine vapor may cause modest increases in impulsive choice 24 hours after exposure (Figure 4).

## 4. Discussion

The present study used a relatively novel rodent e-cigarette delivery system (Qasim et al., 2018) to assess the effects of passive nicotine vapor exposure on serum cotinine levels and impulsive choice. In validation of our model, our findings indicate that passive exposure to nicotine vapor using our rodent vapor exposure system significantly increases serum cotinine levels. Furthermore, our analysis demonstrates that acute passive nicotine vapor exposure can cause short-term increases in impulsive choice in rats. Together, this research not only describes a viable rodent model of human e-cigarette use, but also suggests that exposure to nicotine vapor may affect decision making immediately after exposure, in a similar way to that seen following traditional cigarette use.

Analysis of blood serum cotinine levels immediately following passive exposure to nicotine vapor exposure revealed significantly higher levels of cotinine in the 12 and 24 mg/mL nicotine groups, relative to the 0 mg/mL nicotine group, on all exposure days assessed. Furthermore, cotinine serum levels appeared to increase in the 12 and 24 mg/mL nicotine treatment group between exposure days 1 and 5, and decrease between exposure days 5 and 10. The observed cotinine levels are similar to those seen in another report that examined cotinine levels 7, 10 or 14 days after adolescent male rats were implanted with an osmatic pump that delivered nicotine continuously for 14 days (Torres, Gentil, Natividad, Carcoba, & O’Dell, 2013). When comparing serum cotinine levels between the 24 mg/mL nicotine treatment group and the 12 mg/mL nicotine treatment group, no significant differences were found. However, the 24 mg/mL nicotine group did show a trend for higher serum cotinine levels than the 12 mg/mL on all exposure days assessed. It is also notable that trace amounts of serum cotinine were observed in the 0 mg/mL nicotine group. One previous study exposing rodents to commercially available 0 mg/mL nicotine solutions also reported similarly low amounts of serum cotinine (under 10 ng/mL) and concluded that nicotine contamination had occurred in the 0 mg/mL solution that they used (Smith et al., 2015). We acknowledge that cross contamination may have also occurred in the 0 mg/mL nicotine solution used in this study, but also posit that observed serum cotinine levels in the 0 mg/mL nicotine exposed group could have occurred as a result of third hand nicotine vapor exposure. Third hand exposure has been reported in environments where nicotine vapor is consumed (Bush & Goniewicz, 2015) and despite our efforts to thoroughly clean our chambers after every exposure and use different minitanks and tubing for each nicotine concentration, it may not have been enough to avoid third hand exposure. Future studies investigating the effects of nicotine vapor on the brain and behavior should aim to utilize analytical chemistry techniques to verify the presence of nicotine in commercially available “0 mg/mL” nicotine solutions and could benefit from having completely separate vapor exposure chambers for control groups. Additionally, studies investigating the effects of drugs vapor exposure in rodents would benefit from the development of a delivery system that allows for response contingent vapor administration, as research has shown that passive and self-administration of drugs can have strikingly different effects on behavioral and neurobiological outcomes (Jacobs, Smit, de Vries, & Schoffelmeer, 2003; Metaxas et al., 2010).

When investigating the effects of nicotine vapor on impulsive choice 15-21 days after repeated exposure, no long-term effects were seen in the 12 or 24 mg/mL nicotine vapor exposure groups, when compared to the 0 mg/mL nicotine vapor exposure group. While previous studies examining the effects of nicotine injections on impulsive choice rats did show persistent changes in impulsive choice following 10 day of nicotine vapor exposure, they also demonstrated that after a certain period of abstinence choice preference returned to levels comparable to those seen during control measures (Anderson & Diller, 2010; Dallery & Locey, 2005). Research with humans has also suggested that shifts in choice preference resulting from nicotine exposure may be transient (Bickel et al., 1999; Mitchell, 1999). Therefore, while no long-term effects were seen our study, it is possible that persistent effects of nicotine vapor on choice preference were no longer present when stable performance in the delay discounting task was achieved 15 days after exposure. Future research will need to investigate the effects of repeated nicotine vapor exposure on impulsive choice 2-14 days following treatment.

Analysis of impulsive choice following acute exposure to 24 mg/mL nicotine vapor revealed an immediate shift in choice preference towards the small immediate reward (increase in impulsive choice), relative to choice preference immediately following exposure to 0 mg/mL nicotine vapor, in rats with no history of nicotine vapor exposure (i.e., rats previously exposed to 10 day of 0 mg/mL nicotine vapor). Importantly, post-hoc analysis revealed that the main effect of nicotine vapor on impulsive choice is driven by an increase in impulsivity that is significant only during the 4^th^ block of the task (16 s delay), when optimal choice option is less evident. A similar preference for the larger reward option during block 1 (when there was no delay to the large reward delivery) was observed following acute exposure to both 0 and 24 mg/mL nicotine vapor, suggesting that nicotine vapor exposure does not affect the ability to perceive differences in reward magnitude. Furthermore, a similar preference for the small reward option during block 5 (when the delay to large reward delivery was the longest) was seen following acute exposure to both 0 and 24 mg/mL nicotine vapor, suggesting that nicotine vapor exposure does not affect the ability to detect the temporal delays to large reward delivery and discount the large reward option based on this relatively large associated cost.

The significant effect of acute nicotine vapor exposure on impulsive choice was also seen in rats previously exposed to 10 days of 12 mg/mL nicotine vapor, but not the group that was previously exposed to 10 days of 24 mg/mL nicotine vapor. It is conceivable that a lack of significant acute effects in rats previously exposed to 10 days of 24 mg/mL nicotine vapor is due to an increased tolerance to nicotine, but this notion remains to be investigated. In addition to the immediate increases in impulsive choice observed following exposure to 24 mg/mL nicotine in rats that had 10 days of 12 mg/ml nicotine exposure, a near significant trend for increased impulsive choice was seen in these rats when were tested 24 hours after the acute exposure (Figures 4). This modest increase in impulsive choice 24 hours after a single exposure to 24 mg/mL suggests that significant persistent increases in impulsive choice may be possible if animals were tested 24 hours after repeated exposure to 24 mg/mL nicotine. Taken together, these findings provide important initial evidence that nicotine vapor exposure can indeed increase impulsive choice and highlight the need for future studies to further characterize the effects of nicotine vapor exposure on decision making.

### Summary and conclusions

Traditionally, research has investigated the effects of nicotine on the brain and behavior following exposure to cigarette smoke or nicotine injections. These studies have demonstrated a number of negative health effects, including short and long-term increases in impulsive choice (Bickel et al., 1999; Dallery & Locey, 2005; Locey, Raiff, & Dallery, 2003). Findings from our study demonstrate that exposure to e-cigarette vapor using relatively novel rodent vapor exposure chambers can cause significant short-term, but not long-term, increases in impulsive choice. Future research investigating the effects of nicotine vapor exposure on the brain and behavior will provide much needed information about this novel and largely unknown phenomenon. Relative to traditional cigarettes, e-cigarettes can expose users to different concentrations of nicotine, rates of consumption, and additive chemicals (e.g., propylene glycol, vegetable glycerin), all of which can impact the pulmonary absorption, pharmacodynamics, and health effects of nicotine (Callahan-Lyon, 2014). These differences, along with recent and dramatic increases in e-cigarette use (Gentzke et al., 2019), highlight the need to seek clarity on the effects of nicotine vapor exposure on human health, as well as how the health effects of this novel nicotine delivery system differ from those seen with traditional cigarette use. By doing so, researchers will help address misconceptions regarding the safety of e-cigarettes and improve treatment outcomes for nicotine dependence and e-cigarette related health impairments.

## Acknowledgements

We would like to thank Michelle Martinez, Tania Miramontes, Melissa Ibarra, and Gabriel Frietze for their assistance and technical support on this project.

## References

Anderson, K. G., & Diller, J. W. (2010). Effects of acute and repeated nicotine administration on delay discounting in Lewis and Fischer 344 rats. Behavioural Pharmacology, 21(8), 754–764 710.1097/FBP.1090b1013e328340a328050.

Ascioglu, M., Dolu, N., Golgeli, A., Suer, C., & Ozesmi, C. (2004). Effects of cigarette smoking on cognitive processes. International Journal of Neuroscience, 114, 381–390.

Bickel, W., Odum, A., & Madden, G. (1999). Impulsivity and cigarette smoking: delay discounting in current, never, and ex-smokers. Psychopharmacology, 146, 447–454.

Bush, D., & Goniewicz, M. L. (2015). A pilot study on nicotine residues in houses of electronic cigarette users, tobacco smokers, and non-users of nicotine-containing products. Int J Drug Policy, 26(6), 609–611. doi:10.1016/j.drugpo.2015.03.003

Callahan-Lyon, P. (2014). Electronic cigarettes: human health effects. Tob Control, 23 Suppl 2, ii36–40. doi:10.1136/tobaccocontrol-2013-051470

Chivers, L. L., Hand, D. J., Priest, J. S., & Higgins, S. T. (2016). E-cigarette use among women of reproductive age: Impulsivity, cigarette smoking status, and other risk factors. Prev Med, 92, 126–134. doi:10.1016/j.ypmed.2016.07.029

Chun, L. F., Moazed, F., Calfee, C. S., Matthay, M. A., & Gotts, J. E. (2017). Pulmonary toxicity of e-cigarettes. Am J Physiol Lung Cell Mol Physiol, 313(2), L193–l206. doi:10.1152/ajplung.00071.2017

Dallery, J., & Locey, M. L. (2005). Effects of acute and chronic nicotine on impulsive choice on rats. Behavioural Pharmacology, 16(1), 15–23.

Dani, J. A., & Harris, R. A. (2005). Nicotine addiction and comorbidity with alcohol abuse and mental illness. Nat Neurosci, 8(11), 1465–1470.

Etter, J. F. (2016). A longitudinal study of cotinine in long-term daily users of e-cigarettes. Drug Alcohol Depend, 160, 218–221. doi:10.1016/j.drugalcdep.2016.01.003

Flouris, A. D., Chorti, M. S., Poulianiti, K. P., Jamurtas, A. Z., Kostikas, K., Tzatzarakis, M. N., … Koutedakis, Y. (2013). Acute impact of active and passive electronic cigarette smoking on serum cotinine and lung function. Inhal Toxicol, 25(2), 91–101. doi:10.3109/08958378.2012.758197

Gentzke, A. S., Creamer, M., Cullen, K. A., Ambrose, B. K., Willis, G., Jamal, A., & King, B. A. (2019). Vital Signs: Tobacco Product Use Among Middle and High School Students - United States, 2011-2018. MMWR Morb Mortal Wkly Rep, 68(6), 157–164. doi:10.15585/mmwr.mm6806e1

Gilpin, N. W., Whitaker, A. M., Baynes, B., Abdel, A. Y., Weil, M. T., & George, O. (2014). Nicotine vapor inhalation escalates nicotine self-administration. Addict Biol, 19(4), 587–592. doi:10.1111/adb.12021

Hall, B. J., Wells, C., Allenby, C., Lin, M. Y., Hao, I., Marshall, L., … Levin, E. D. (2014). Differential effects of non-nicotine tobacco constituent compounds on nicotine self-administration in rats. Pharmacol Biochem Behav, 120, 103–108. doi:10.1016/j.pbb.2014.02.011

Jacobs, E. H., Smit, A. B., de Vries, T. J., & Schoffelmeer, A. N. (2003). Neuroadaptive effects of active versus passive drug administration in addiction research. Trends Pharmacol Sci, 24(11), 566–573. doi:10.1016/j.tips.2003.09.006

Javadi-Paydar, M., Kerr, T. M., Harvey, E. L., Cole, M., & Taffe, M. A. (2019). Effects of nicotine and THC vapor inhalation administered by an electronic nicotine delivery system (ENDS) in male rats. Drug Alcohol Depend, 198, 54–62. doi:10.1016/j.drugalcdep.2019.01.027

Locey, M., Raiff, B., & Dallery, J. (2003). Effects of nicotine on impulsive choice in rats. Paper presented at the Annual Meeting of the Association for Behavior Analysis, San Francisco.

Lozano, P., Barrientos-Gutierrez, I., Arillo-Santillan, E., Morello, P., Mejia, R., Sargent, J. D., & Thrasher, J. F. (2017). A longitudinal study of electronic cigarette use and onset of conventional cigarette smoking and marijuana use among Mexican adolescents. Drug Alcohol Depend, 180, 427–430. doi:10.1016/j.drugalcdep.2017.09.001

Mansvleder, H. D., & McGehee, D. S. (2002). Cellular and synaptic mechanisms of nicotine addiction. Journal of Neurobiology, 53, 606–617.

Mendez, I. A., Gilbert, R. J., Bizon, J. L., & Setlow, B. (2012). Effects of acute administration of nicotinic and muscarinic cholinergic agonists and antagonists on performance in different cost-benefit decision making tasks in rats. Psychopharmacology.

Mendez, I. A., Simon, N. W., Hart, N., Mitchell, M. R., Nation, J. R., Wellman, P. J., & Setlow, B. (2010). Self-administered cocaine causes long-lasting increases in impulsive choice in a delay discounting task. Behavioral Neuroscience, 124(4), 470–477.

Metaxas, A., Bailey, A., Barbano, M. F., Galeote, L., Maldonado, R., & Kitchen, I. (2010). Differential region-specific regulation of alpha4beta2* nAChRs by self-administered and non-contingent nicotine in C57BL/6J mice. Addict Biol, 15(4), 464–479. doi:10.1111/j.1369-1600.2010.00246.x

Mitchell, S. H. (1999). Measures of impulsivity in cigarette smokers and nonsmokers. Psychopharmacology, 146, 455–464.

Mitchell, S. H. (2004). Measuring impulsivity and modeling its association with cigarette smoking. Behavioral and Cognitive Neuroscience Reviews, 3(5), 261–275.

Qasim, H., Karim, Z. A., Silva-Espinoza, J. C., Khasawneh, F. T., Rivera, J. O., Ellis, C. C., … Alshbool, F. Z. (2018). Short-Term E-Cigarette Exposure Increases the Risk of Thrombogenesis and Enhances Platelet Function in Mice. J Am Heart Assoc, 7(15). doi:10.1161/jaha.118.009264

Sengupta, P. (2013). The Laboratory Rat: Relating Its Age With Human’s. Int J Prev Med, 4(6), 624–630.

Simon, N. W., Mendez, I. A., & Setlow, B. (2007). Cocaine exposure causes long-term increases in impulsive choice. Behavioral Neuroscience, 121(3), 543–549.

Smith, D., Aherrera, A., Lopez, A., Neptune, E., Winickoff, J. P., Klein, J. D., … McGrath-Morrow, S. A. (2015). Adult Behavior in Male Mice Exposed to E-Cigarette Nicotine Vapors during Late Prenatal and Early Postnatal Life. PLoS One, 10(9), e0137953. doi:10.1371/journal.pone.0137953

Torres, O. V., Gentil, L. G., Natividad, L. A., Carcoba, L. M., & O’Dell, L. E. (2013). Behavioral, Biochemical, and Molecular Indices of Stress are Enhanced in Female Versus Male Rats Experiencing Nicotine Withdrawal. Front Psychiatry, 4, 38. doi:10.3389/fpsyt.2013.00038

Yuan, M., Cross, S. J., Loughlin, S. E., & Leslie, F. M. (2015). Nicotine and the adolescent brain. J Physiol, 593(Pt 16), 3397–3412. doi:10.1113/jp270492

